## Supplementary Figures and Methods for "Robust analysis of allele-specific copy number alterations from scRNA-seq data with XClone"

[1]School of Biomedical Sciences, The University of Hong Kong, Hong Kong SAR, China
[2]Center for Translational Stem Cell Biology, Hong Kong Science and Technology Park, Hong Kong SAR, China
[3]Department of Pathology, School of Clinical Medicine, LKS Faculty of Medicine, The University of Hong Kong, Queen Mary Hospital, Hong Kong SAR, China
[4]The Jockey Club Centre for Clinical Innovation and Discovery, LKS Faculty of Medicine, The University of Hong Kong, Pokfulam, Hong Kong SAR, China
[5]Centre for PanorOmic Sciences, LKS Faculty of Medicine, The University of Hong Kong, Pokfulam, Hong Kong SAR, China
[6]Division of Computational Genomics and Systems Genetics, German Cancer Research Center (DKFZ), Heidelberg, Germany
[7]Genome Biology Unit, European Molecular Biology Laboratory, Heidelberg, Germany
[8]Department of Statistics and Actuarial Science, The University of Hong Kong, Pokfulam, Hong Kong SAR, China
[*]Co-first authors

#### Supplementary Tables

See the full Supplementary Tables in a separate Excel file, including:

**Supplementary Table S1**: Tumor single cell transcriptomic dataset information and CNA profiles.

**Supplementary Table S2**: BCH869 CNA ground truth.

**Supplementary Table S3**: Number of consensus cells and genes of five methods for benchmarking on BCH869 scRNA-seq datasets.

**Supplementary Table S4**: Running time and memory usage of five methods on three scRNA-seq datasets.

**Supplementary Table S5**: Summarized benchmarking utilities of five methods.

**Supplementary Table S6**: SNPs within significant DEGs on chromosome 14 in the BCH869 dataset.

**Supplementary Table S7**: GBM 2R Clone1 CNA ground truth.

**Supplementary Table S8**: Number of consensus cells and genes of five methods for benchmarking on 2R Clone 1 in GBM snRNA-seq dataset.

**Supplementary Table S9**: GX109-T1c CNA ground truth.

**Supplementary Table S10**: CNA ground truth for the simulation of allelic copy loss in GX109-T1c data.

**Supplementary Table S11**: Cell annotations for the simulation of allelic copy loss in GX109-T1c data.

**Supplementary Table S12**: Comprehensive benchmarking performance on simulated datasets.

### Supplementary Figures

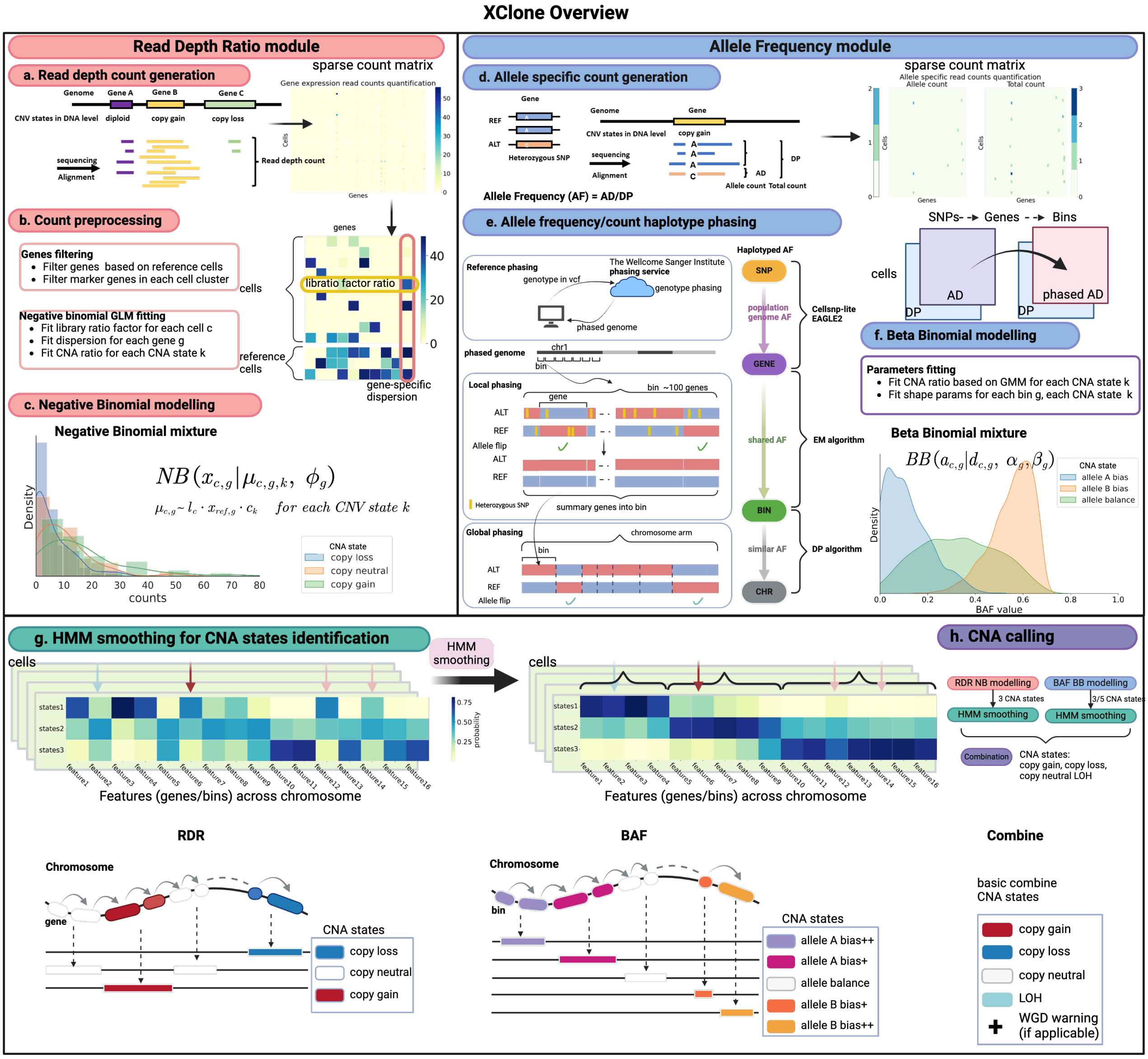


Figure S1: Overview of XClone algorithm of two independent modules: Read Depth Ratio (RDR) module and B-allele frequency (BAF) module. For RDR module (a-c), (a) generation of the read counts in a cell by gene matrix. (b) Preprocessing of genes quality control and Negative binomial GLM parameters fitting with the reference cells. (c) Negative binomial mixture modelling and cell-by-gene-by-state likelihood tensor can be computed for this module. For BAF module(d-f), (d) generation of the allele-specific read counts in a cell by gene matrix with heterozygous SNPs. (e) 3 steps of B-allele frequency haplotype phasing: Reference phasing, local phasing and global phasing. (f) Beta binomial modelling and cell-by-gene_bin-by-state likelihood tensor can be computed for this module. (g) A hidden Markov model and its forward-backward algorithm to smooth the assignment probabilities along genomic coordinates. The lower panel illustrates the transition of CNA states via HMM modelling within the RDR and BAF modules, respectively. (h) CNA calling with each module and combined mode,basic CNA states legend in the final combination(combination strategy shown in Supp. Fig. S2).

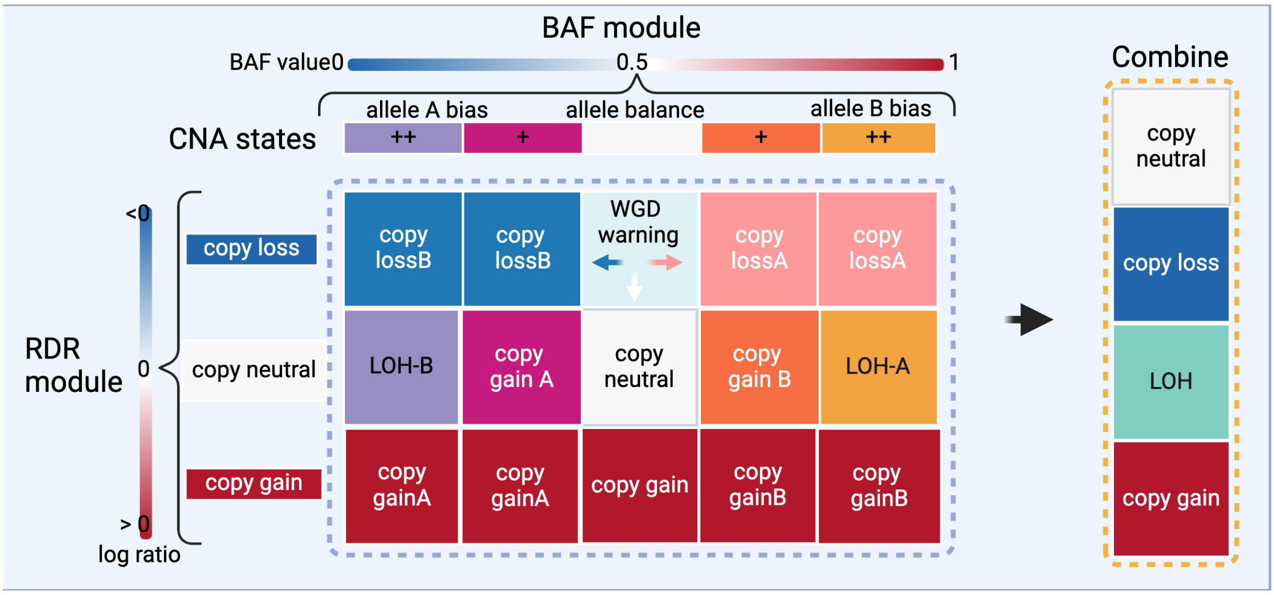


Figure S2: XClone combination strategy: 3-by-5 state probability matrix between RDR and BAF modules will be further aggregated into 6 allelic-specific states: copy neutral, gain, loss A, loss B, LoH A and LoH B.

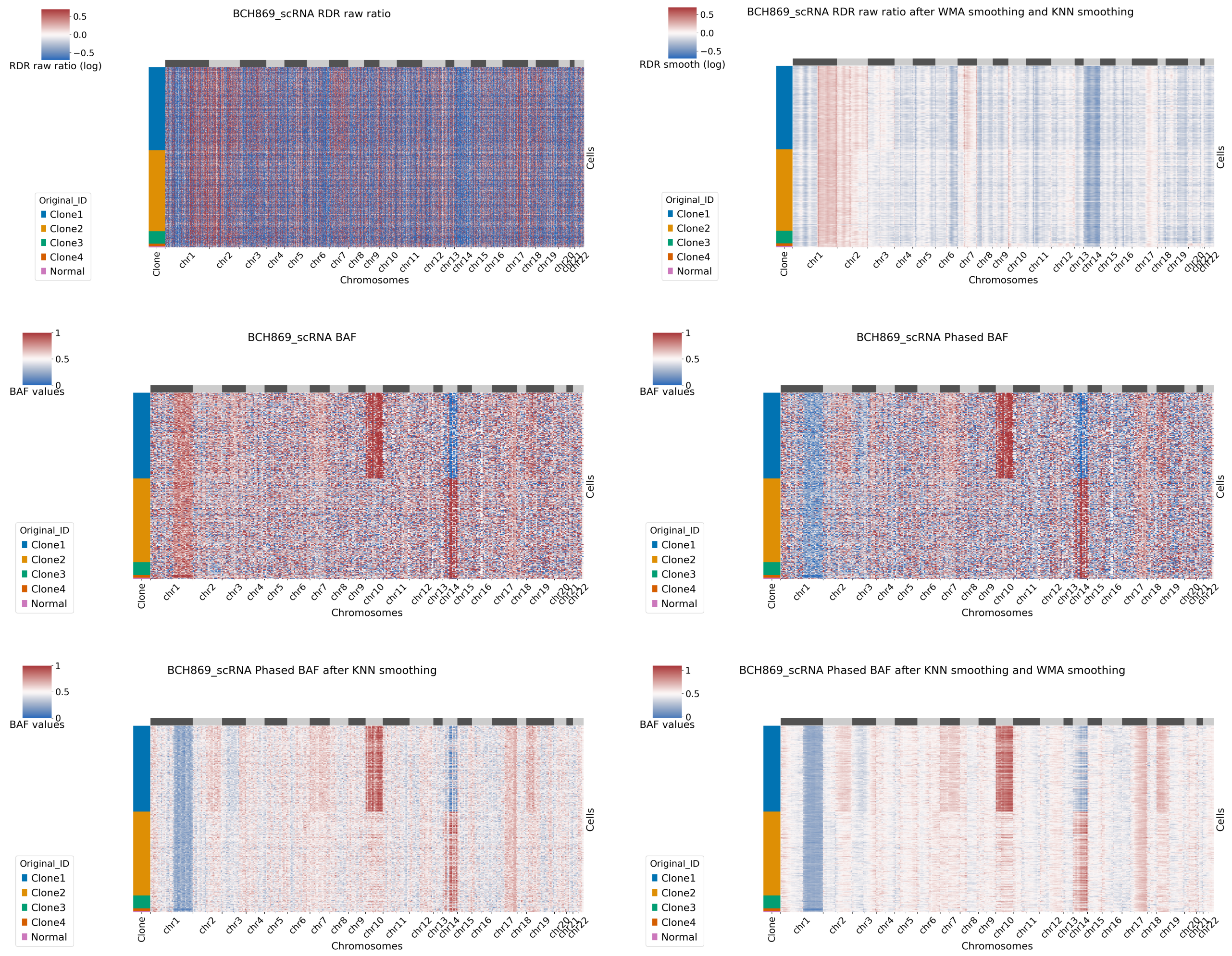


Figure S3: Heatmaps of BCH869 scRNA-seq raw read depth ratio (RDR) and B Allele Frequency (BAF) before and after smoothing generated by XClone..

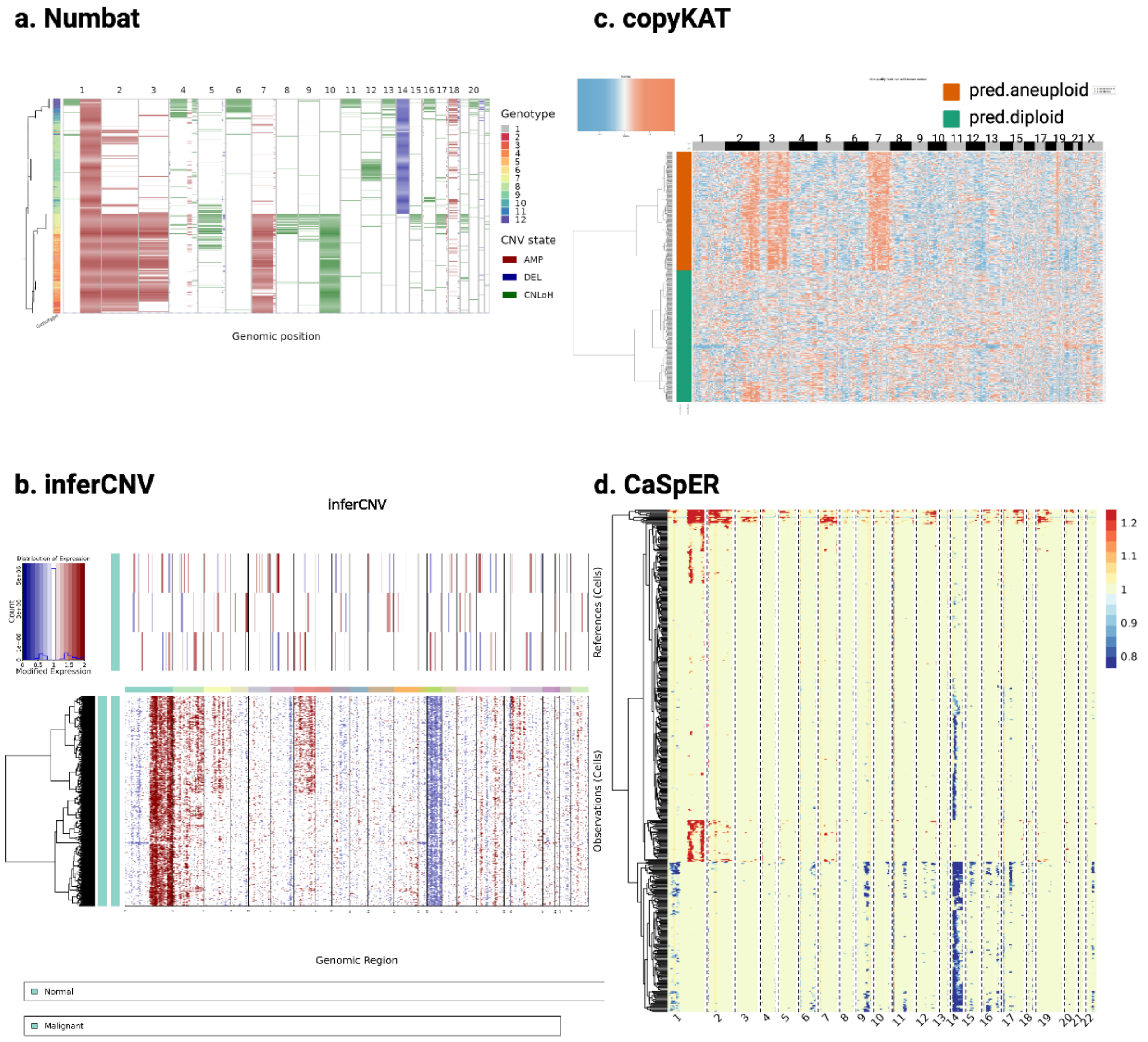


Figure S4: Heatmaps of BCH869 scRNA-seq generated by the four tools for benchmarking. (a) Numbat (Same with Fig. 2e). (b) InferCNV with cell annotation labels of Malignant and Normal. (c) CopyKAT. (d) CaSpER, all tools use 3 normal cells as reference.

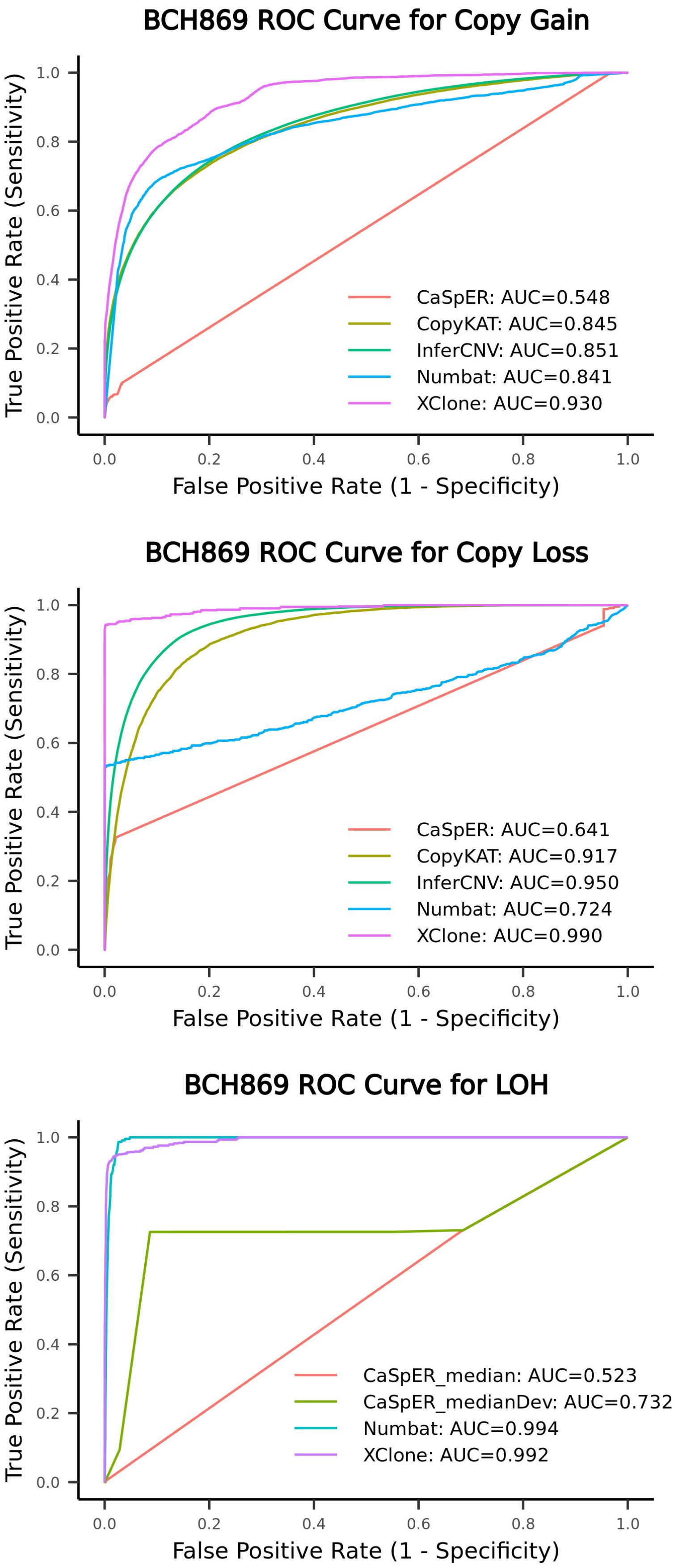


Figure S5: Assessment of performance in identification of copy number gain, copy number loss and loss of heterozygosity on BCH869 (at chromosome arm scale).

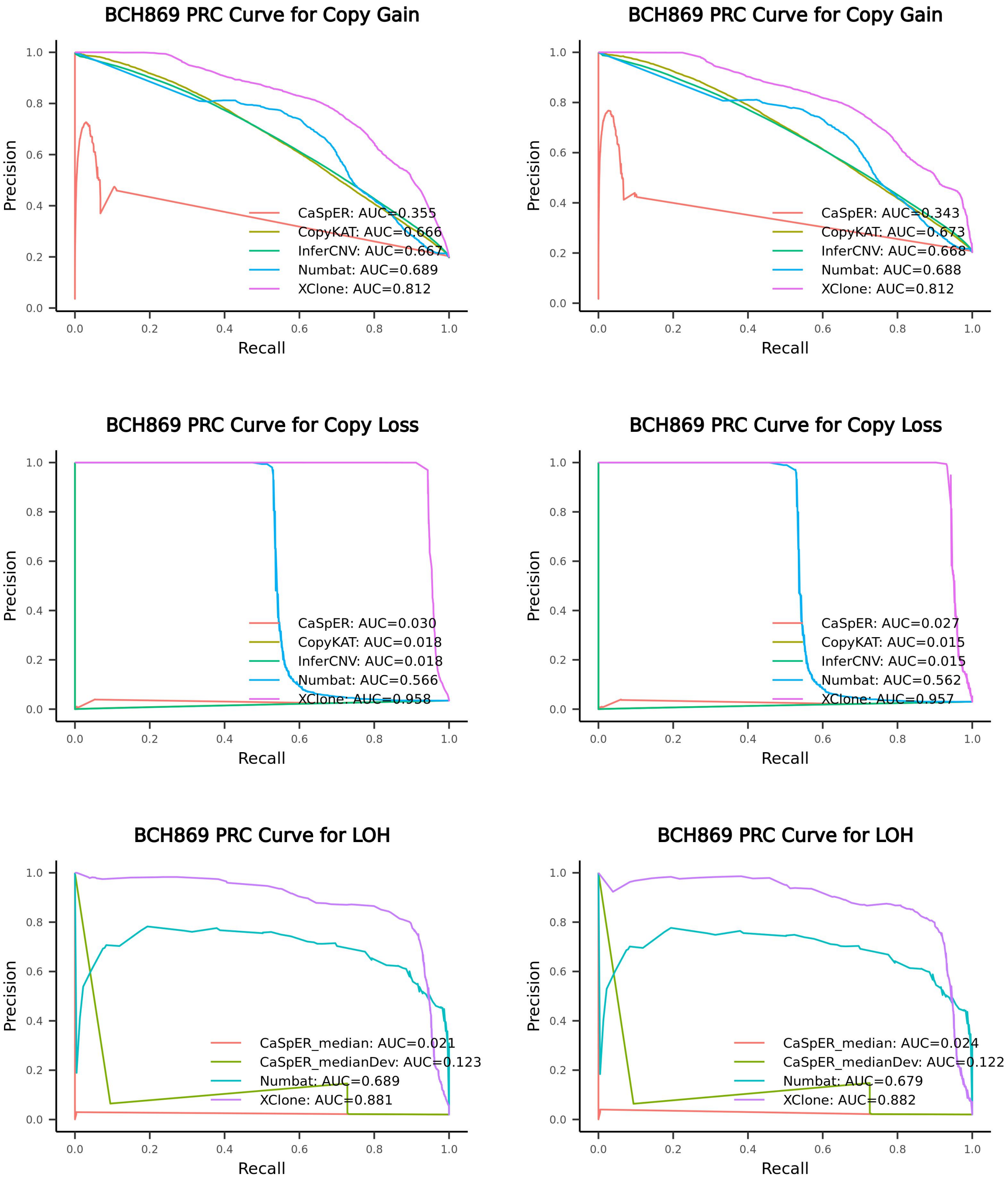


Figure S6: Assessment of performance (PRC curve) in the identification of copy number gain, copy number loss and loss of heterozygosity on BCH869 (left panel: gene scale; right panel: chromosome arm scale).

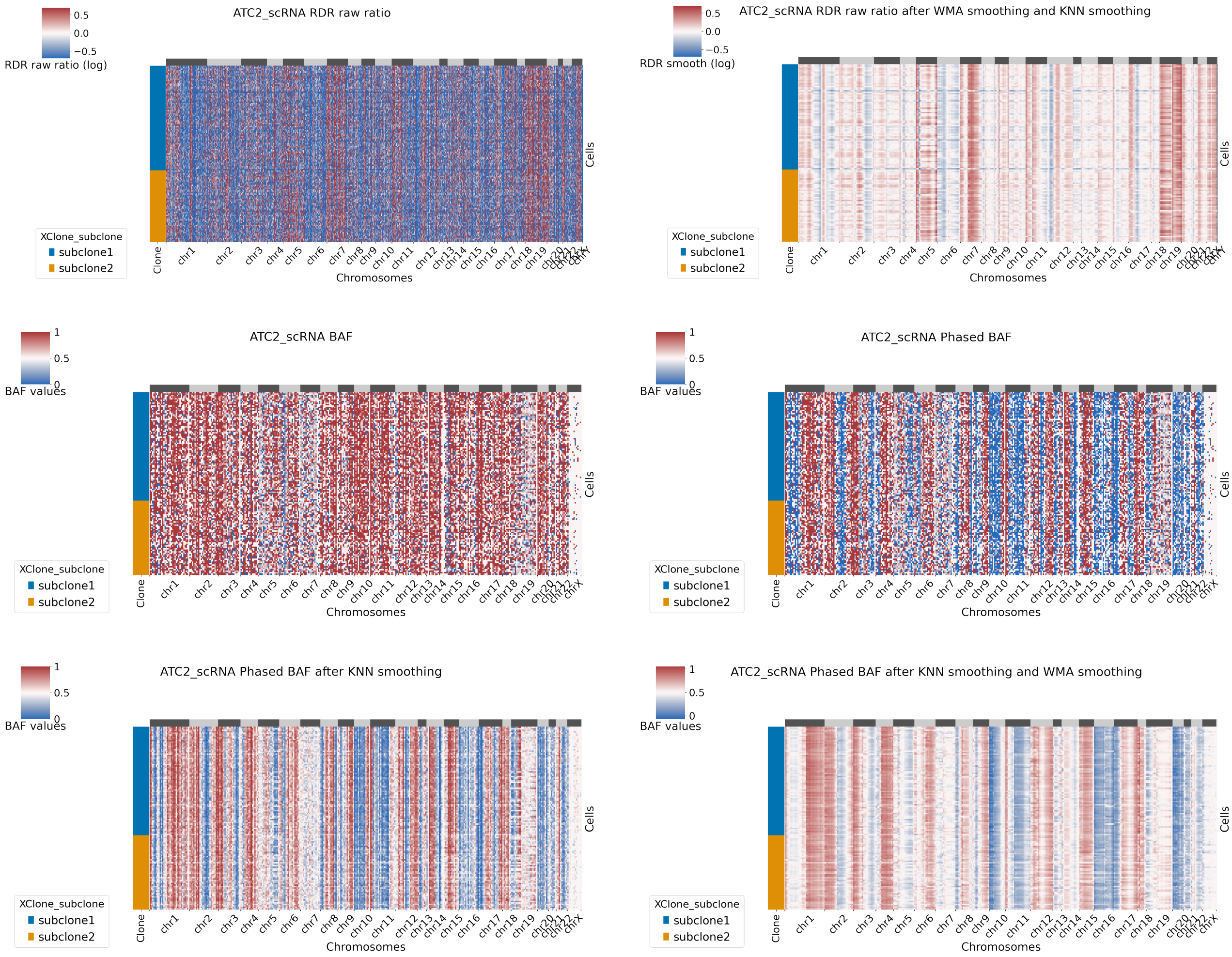


Figure S7: Heatmaps of ATC2 scRNA-seq (tumor subclone) raw read depth ratio (RDR) and B Allele Frequency (BAF) before and after smoothing generated by XClone.

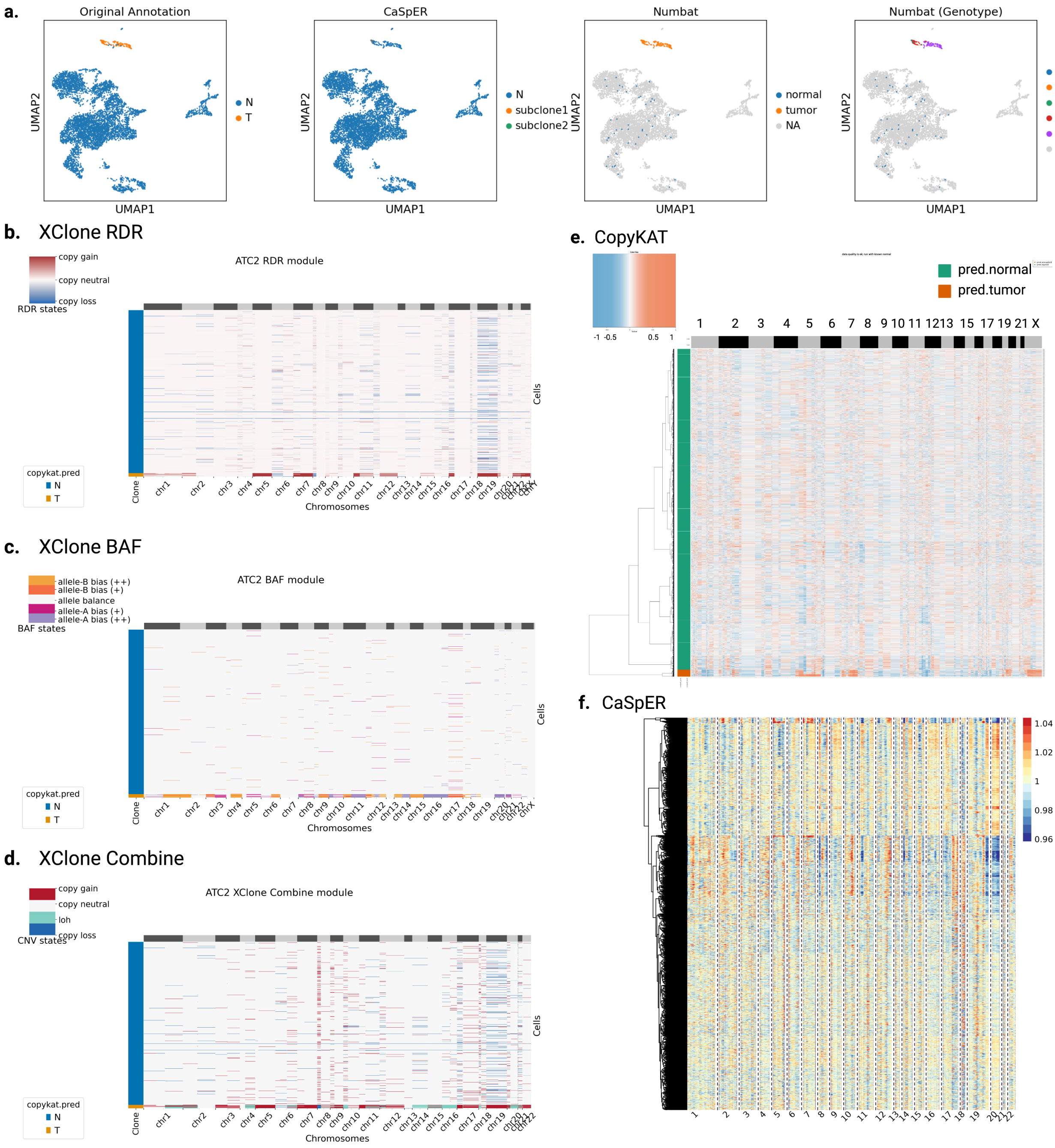


Figure S8: ATC2 scRNA-seq sample. (a) UMAP of ATC2 with different types of cell annotation: original cell annotation (Tumor/Normal) provided by CopyKAT, tumor subclones identified by CaSpER and Numbat. (b-d) Heatmap of ATC2 sample CNA identification results by XClone RDR module, BAF module and combined module for all cells. (e) Heatmap of ATC2 sample CNA identification results by CopyKAT with Normal cells as reference. (f) Heatmap of ATC2 sample CNA identification results by CaSpER with Normal cells as reference.

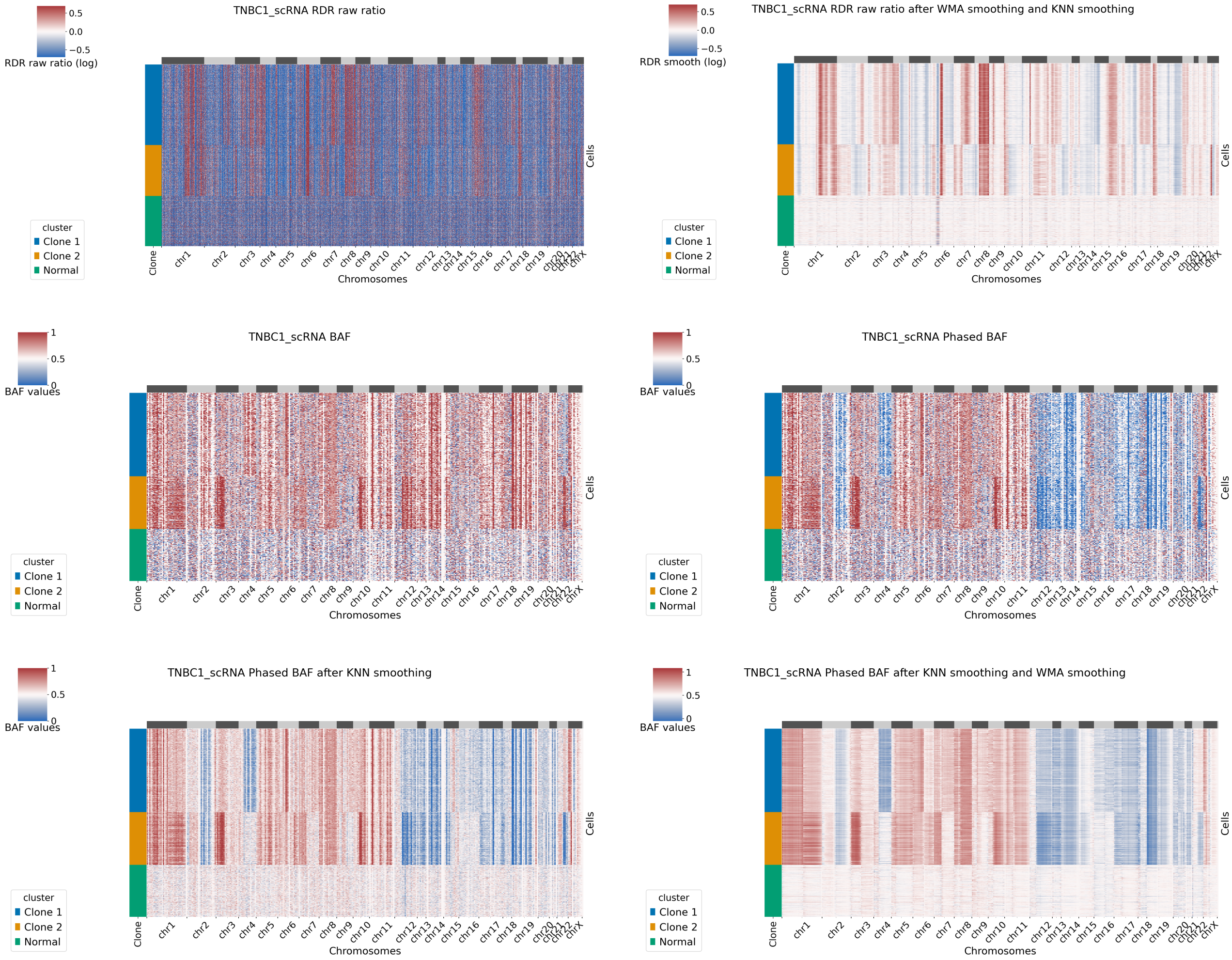


Figure S9: Heatmaps of TNBC1 scRNA-seq raw read depth ratio (RDR) and B Allele Frequency (BAF) before and after smoothing generated by XClone.

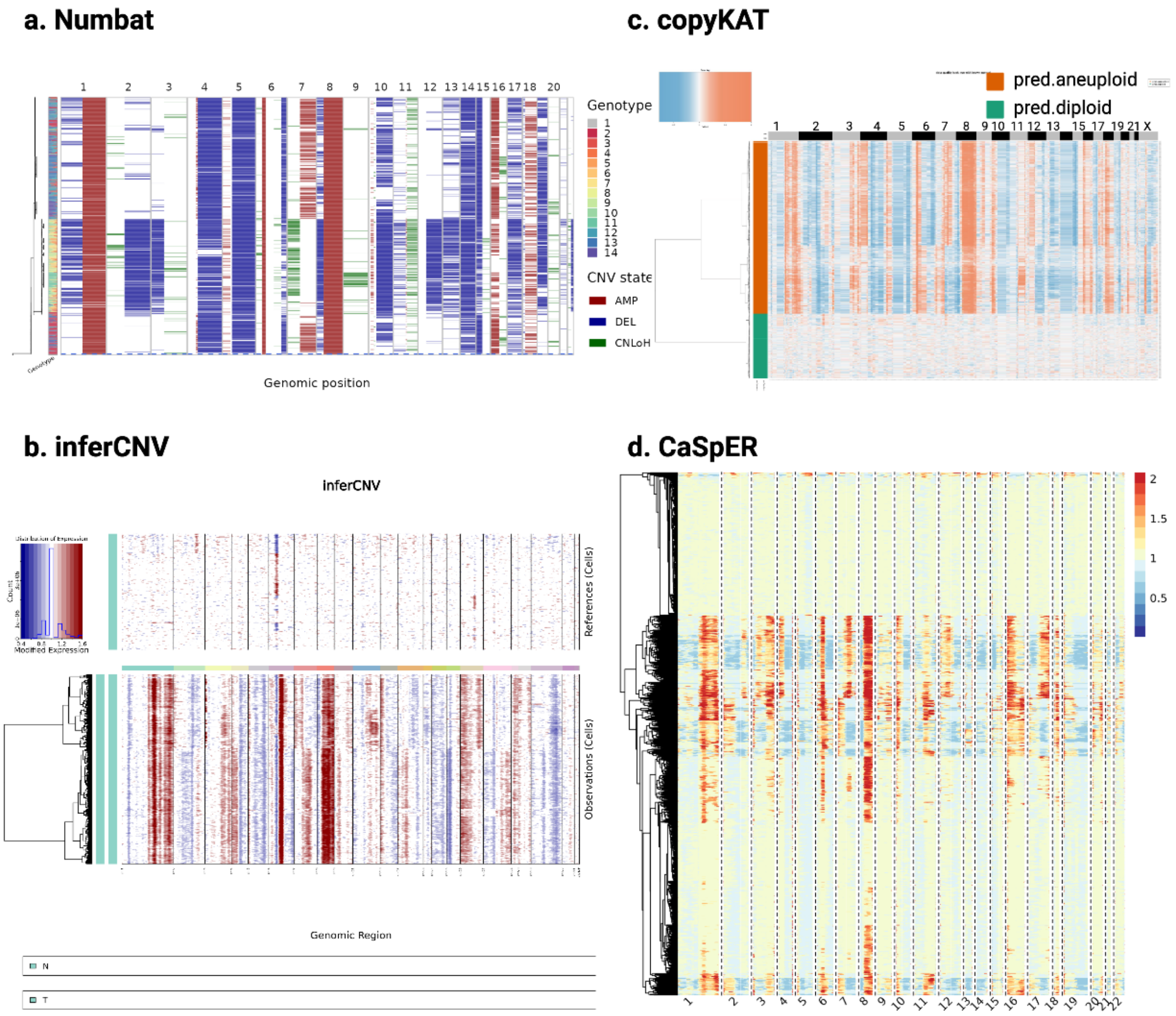


Figure S10: Heatmaps of TNBC1 scRNA-seq generated by the three tools for comparison. (a) Numbat (Same with Fig. 4f) (b) InferCNV (c) CopyKAT. (d) CaSpER with Normal cells as reference.

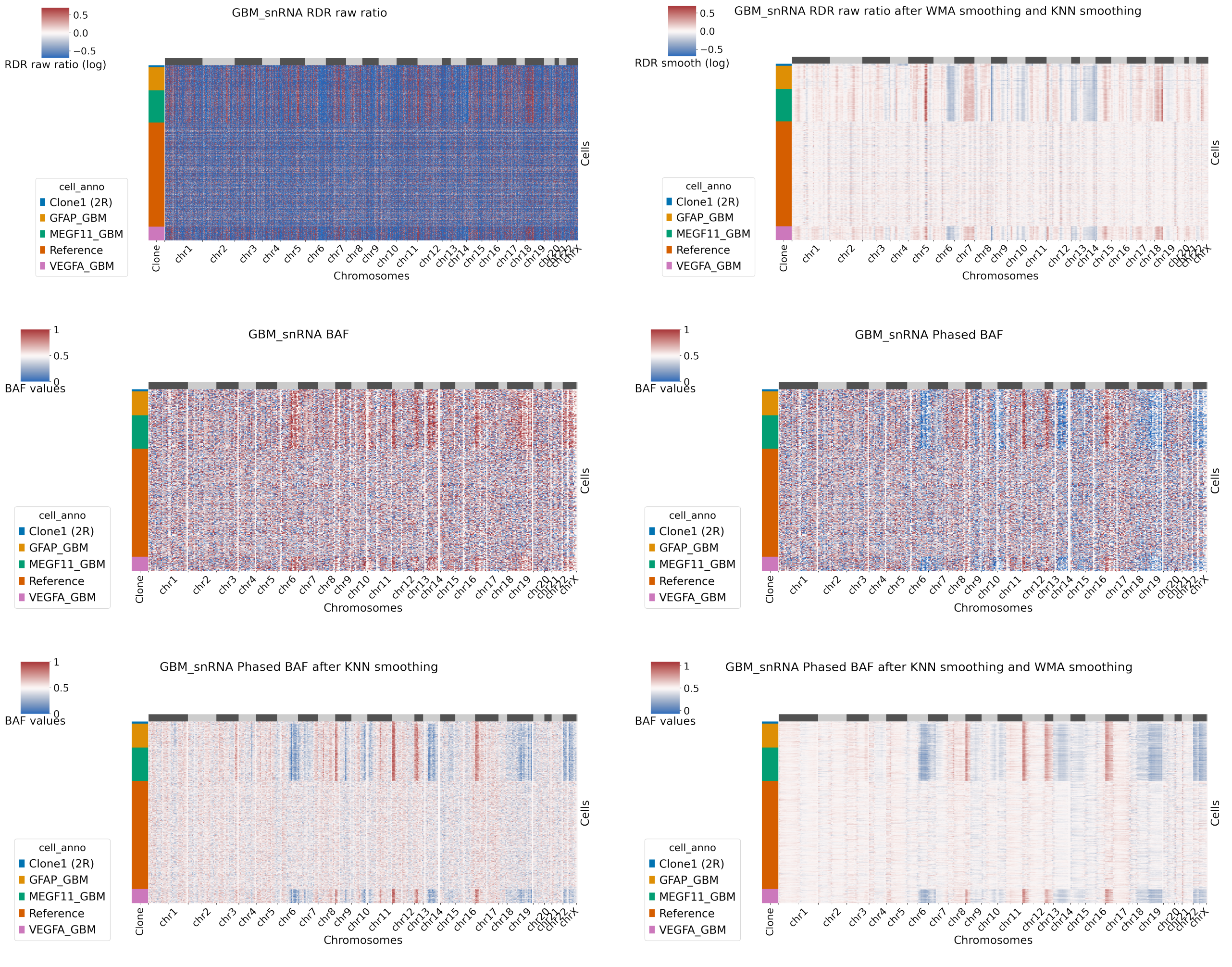


Figure S11: Heatmaps of GBM snRNA-seq raw read depth ratio (RDR) and B Allele Frequency (BAF) before and after smoothing generated by XClone.

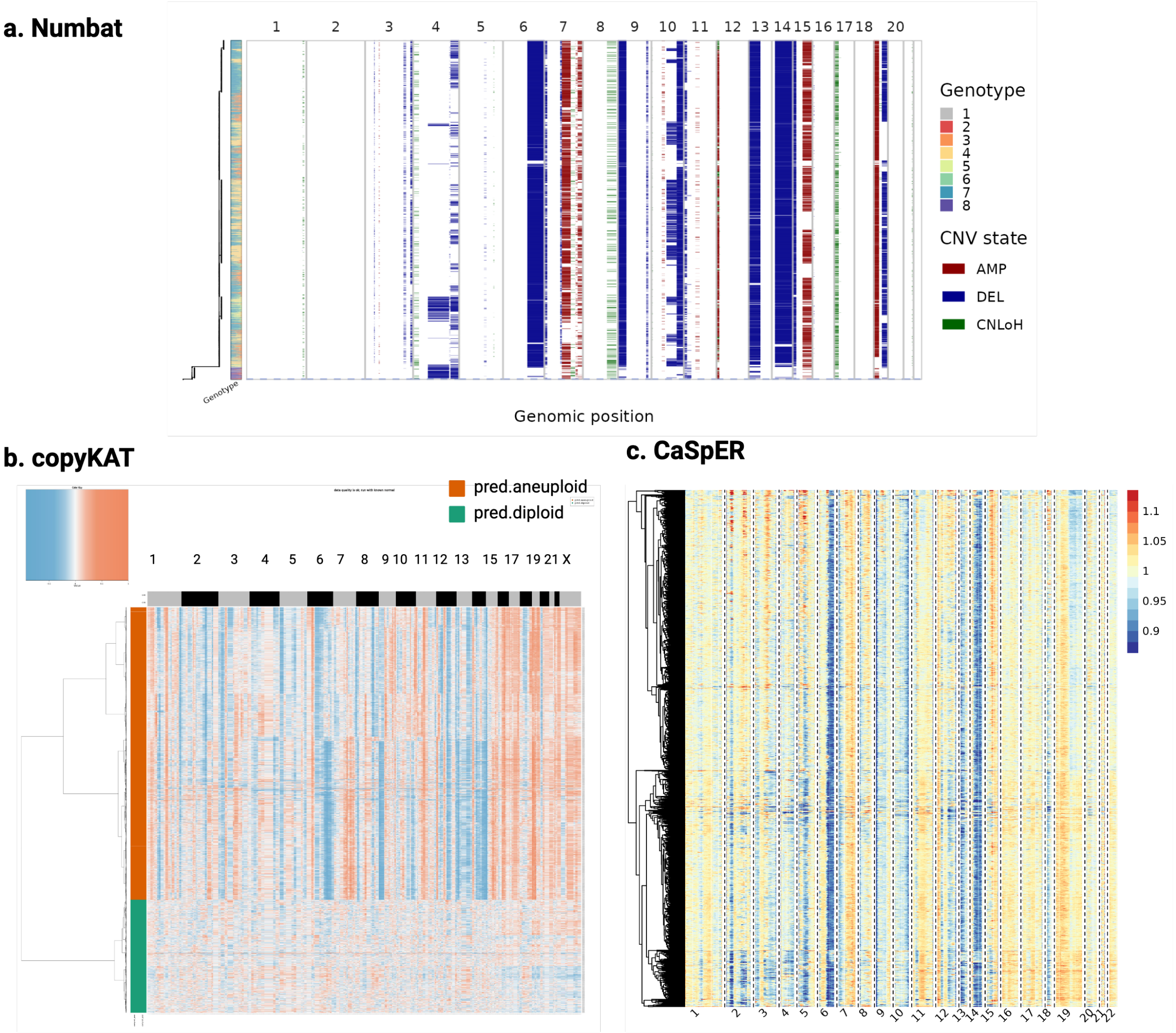


Figure S12: Heatmaps of GBM scRNA-seq generated by the three tools for comparison. (a) Numbat (b) CopyKAT. (c) CaSpER. Note: Numbat and CaSpER outputs exclude the reference cells.

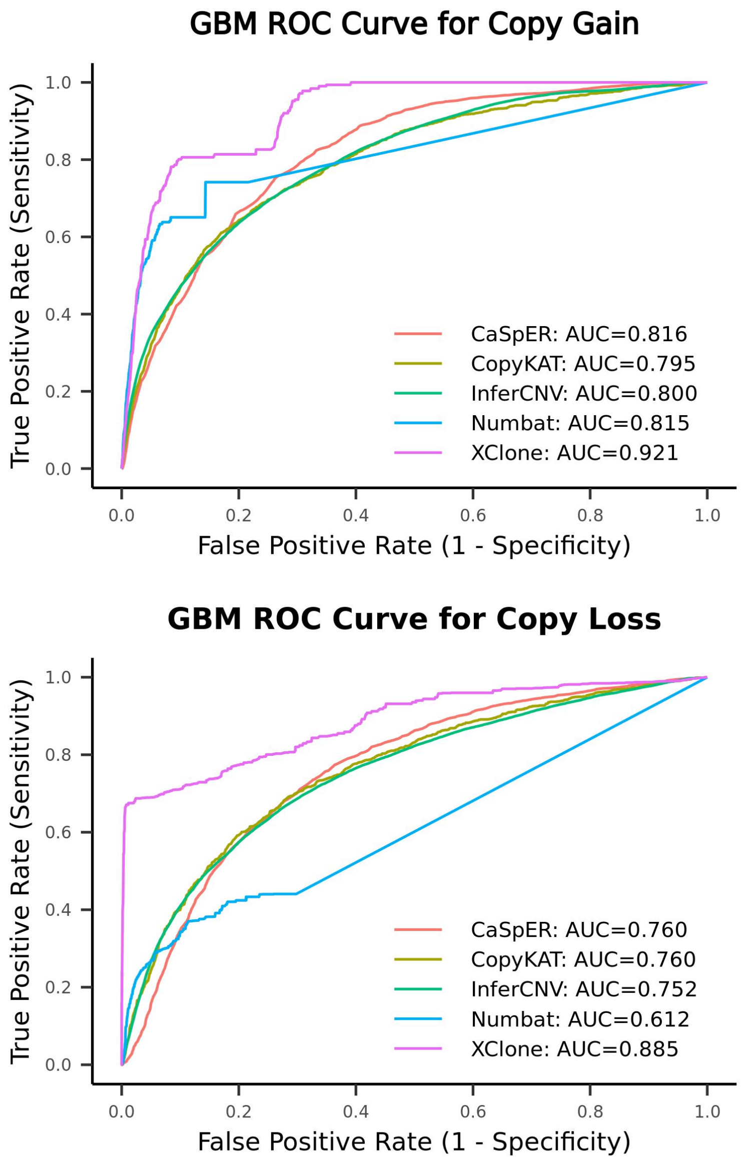


Figure S13: Assessment of performance in identification of copy number gain, copy number loss on GBM dataset (at chromosome arm scale).

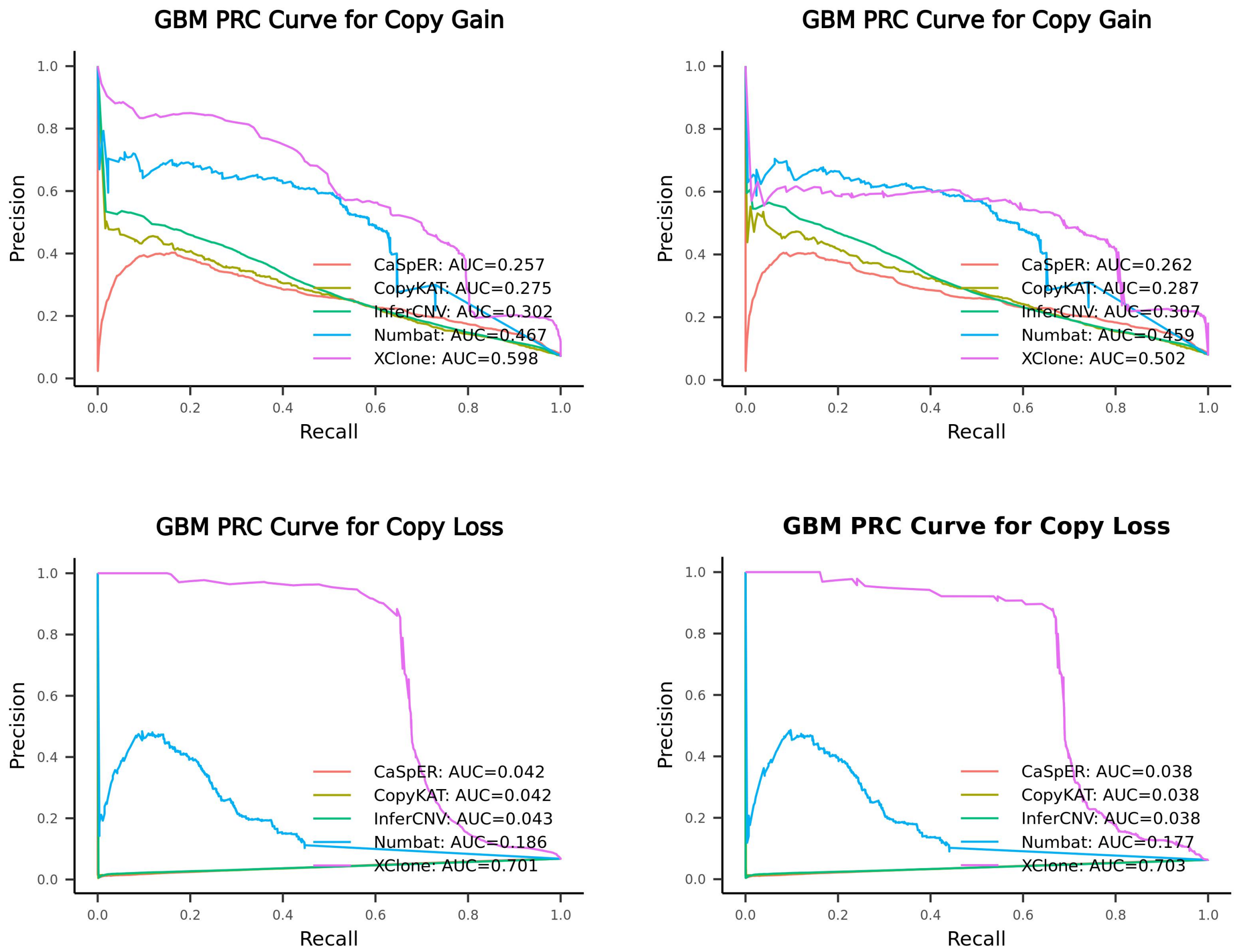


Figure S14: Assessment of performance (PRC curve) in the identification of copy number gain, copy number loss on GBM (left panel: gene scale; right panel: chromosome arm scale).

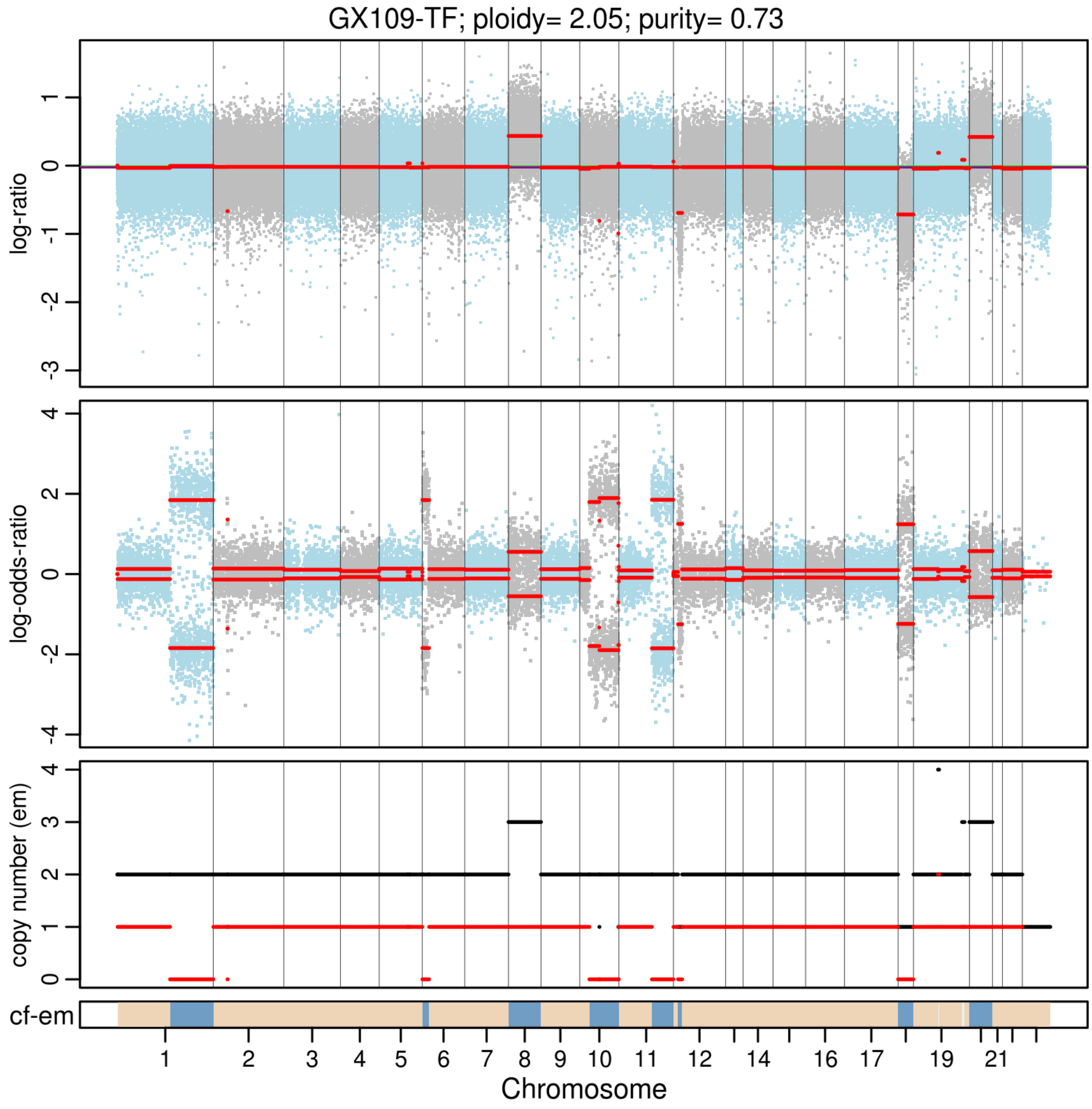


Figure S15: Bulk exome-sequencing of gastric cancer tissue (GX109-TF, taken adjacent to the tissue block used for sc-RNAseq and sc-DNAseq) delineates regions of copy number gain, copy number loss and loss of heterozygosity. Copy number inference was performed using patient's own blood DNA as normal reference, and detected using CNV-facet.

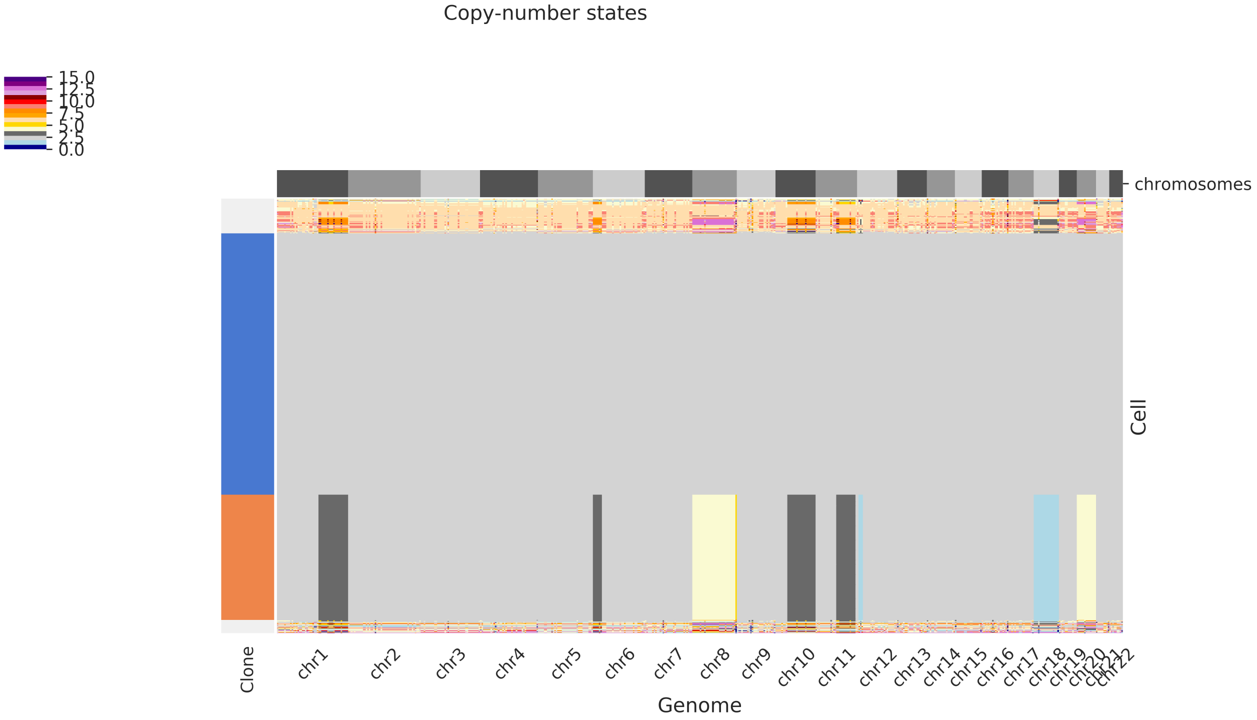


Figure S16: Copy number inference performed on GX109-T1c scDNA-seq using CHISEL Method, which delineates regions of copy number gain (light yellow), copy number loss (light blue) and loss of heterozygosity(dark grey).

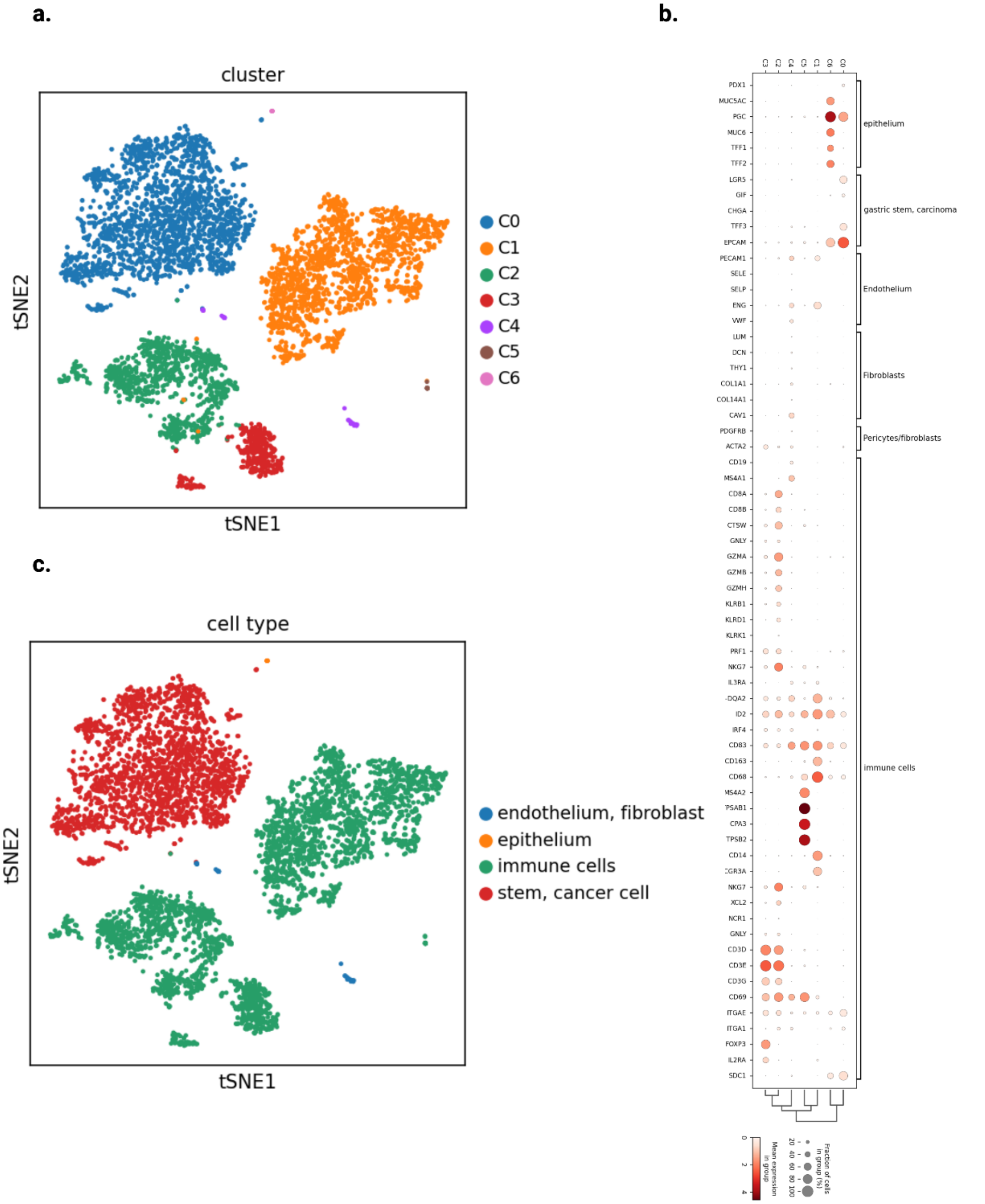


Figure S17: GX109-T1c scRNA-seq manually annotation. (a) TSNE plot by Seurat (7 clusters: C0-C6) at the resolution of 0.1 in the FindClusters Function. (b) Dotplot shows the celltype gene markers pattern. (c) We annotate the C0 as Stem, Cancer cell, C6 as epithelium, C1-C3 as immune cells and C4 as endothelium or fibroblast cells.

GX109-T1c Simulation: Ground truth

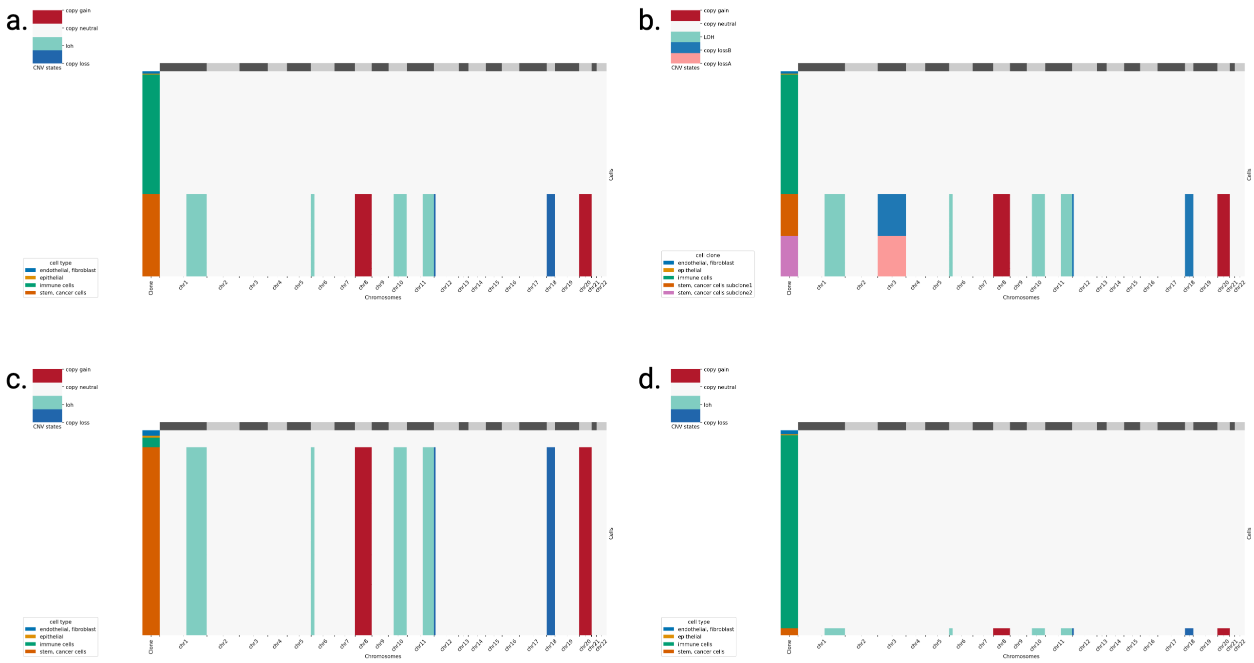

Figure S18: Illustration of the seed data and CNA ground truth for the three simulation scenarios. (a) The CNA profile of the seed dataset GX109-T1c. (b) Ground truth for the simulation of allele-specific copy number loss on chromosome 3 in different sub-tumor clones. This figure is identical to the main Figure 6c for easier reference. (c) Ground truth for the simulation of reference cells downsampling to minor quantities. (d) Ground truth for the simulation of tumor cell downsampling to limited quantities.

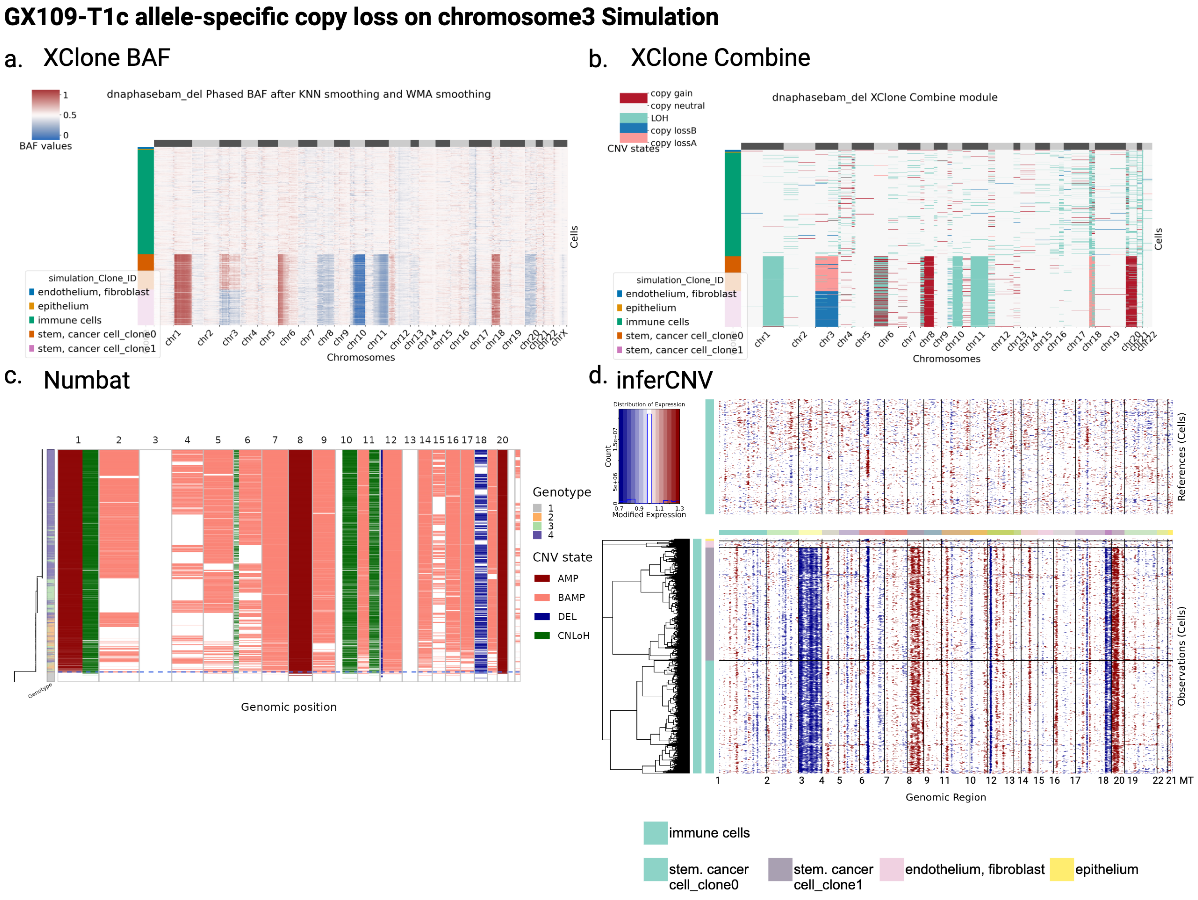


Figure S19: Comparative heatmap results by 3 tools, XClone, Numbat and inferCNV, on the simulated data in scenario 1 with clone specific allele loss in chr3 (ground truth shown in Supp. Fig. S18b). (a-b) XClone shows good performance on the simulated allele-specific copy loss in chromosome 3, with visualization of the smoothed BAF signal (a) and called CNA states (b). (c-d) Numbat and inferCNV cannot show the expected CNA profile well on the simulated allele-specific copy loss in chromosome 3. (c) Numbat cannot call the copy loss in chromosome 3, not to mention the allele specific situation. (d) InferCNV can call the copy loss in chromosome 3 but no allele specific information and no LOH events were detected as expected. It should be noted that the outputs from Numbat do not include the reference cells.

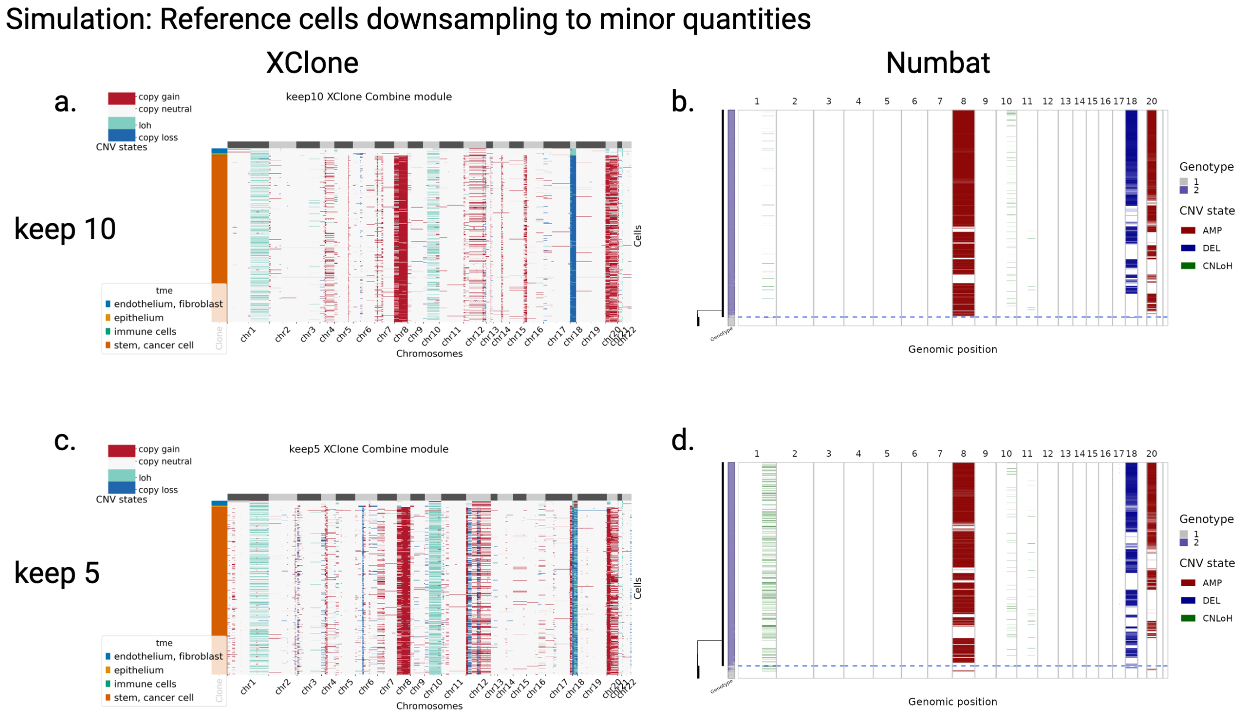


Figure S20: Comparative heatmap results by two tools, Numbat and XClone, on the simulated data in scenario 2 with reference cells downsampling to minor quantities (ground truth shown in Supp. Fig. S18c). (a-b) Keeping only 10 reference cells in the simulated dataset, results from XClone and Numbat, respectively. Similarly, (c-d) are heatmaps for keeping 5 reference cells in the simulated dataset from XClone and Numbat, respectively. It should be noted that the outputs from Numbat do not include the reference cells.

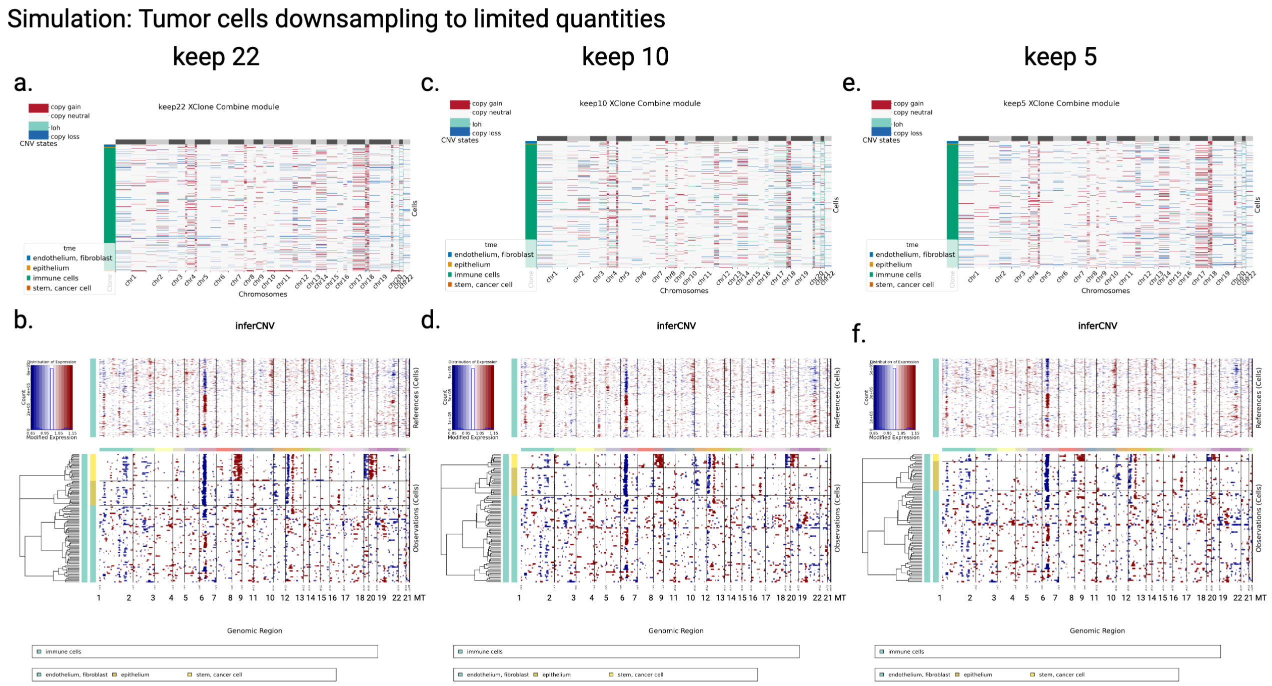


Figure S21: Comparative heatmap results by two tools, XClone and InferCNV, on the simulated data in scenario 2 with tumor cell downsampling to limited quantities (ground truth shown in Supp. Fig. S18d). (a-b) Keeping only 22 tumor cells in the simulated dataset, results from XClone and InferCNV, respectively. Similarly, (c-d) and (e-f) are for keeping 10 tumor cells and 5 tumor cells in the simulated dataset, respectively.

#### Supplementary Algorithms

**Algorithm 1:** EM algorithm for local phasing over $G$ genes in a gene_bin

**Input** allelic count matrices $A$ and $B$ (shape $C$-by-$G$) for $C$ cells and $G$ genes
**Initialize** allele frequency $\boldsymbol{\rho}_{1:C}$ for $C$ cells, flipping probability $\boldsymbol{z}_{1:G}$ for $G$ genes
**while** *not converged* **do**
  **E step**: With current $\boldsymbol{\rho}$, calculate flipping probability for each gene:
  $\boldsymbol{z}_{1:G} = \frac{\exp\{\log(\boldsymbol{\rho})A + \log(1-\boldsymbol{\rho})B\}}{\exp\{\log(\boldsymbol{\rho})A + \log(1-\boldsymbol{\rho})B\} + \exp\{\log(1-\boldsymbol{\rho})A + \log(\boldsymbol{\rho})B\}}$.
  **M step**: Maximizing likelihood on parameters with current expectation $\boldsymbol{z}$:
  $\boldsymbol{\rho}_{1:C} = \frac{A \times \boldsymbol{z} + B \times (\mathbf{1} - \boldsymbol{z})}{(A+B) \times \mathbf{1}}$,
  $\mathcal{L}(\boldsymbol{\rho}) = \sum \log\{\exp\{\log(\boldsymbol{\rho})A + \log(1-\boldsymbol{\rho})B\} + \exp\{\log(1-\boldsymbol{\rho})A + \log(\boldsymbol{\rho})B\}\}$.
  **Update** $\log \mathcal{L}(\boldsymbol{\rho})$ and check convergence
**return** $\boldsymbol{\rho}, \boldsymbol{z}, \log \mathcal{L}(\boldsymbol{\rho})$

**Algorithm 2:** Dynamic programming for global phasing a chromosome arm

**Input:** allele frequency matrix $\boldsymbol{\rho}$ (shape $C$-by-$K$) for $C$ cells and $K$ gene_bins
**Output:** $\boldsymbol{z}, \boldsymbol{\rho}$ for flipping state $\boldsymbol{z}_{1:K}$ and phased allele frequency matrix $\boldsymbol{\rho}_{1:C,1:K}$
**Function** `GlobalPhasing`($\boldsymbol{\rho}$):
  **if** $K == 1$ **then**
    $\boldsymbol{z} = \{0\}$ // no flipping
    **return** $\boldsymbol{z}, \boldsymbol{\rho}$
  **else**
    $\boldsymbol{z}_{1:K-1}, \boldsymbol{\rho}_{1:C,1:K-1} =$ `GlobalPhasing`($\boldsymbol{\rho}_{:,1:K-1}$)
    $d_0 = ||\boldsymbol{\rho}_{:,K} - \boldsymbol{\rho}_{:,K-1}||_2$ // Euclidean distance
    $d_1 = ||(\mathbf{1} - \boldsymbol{\rho}_{:,K}) - \boldsymbol{\rho}_{:,K-1}||_2$
    **if** $d_0 < d_1$ **then**
      $z_K = 0$
      $\boldsymbol{\rho}_{:,K} = \boldsymbol{\rho}_{:,K}$
    **else**
      $z_K = 1$
      $\boldsymbol{\rho}_{:,K} = \mathbf{1} - \boldsymbol{\rho}_{:,K}$
    **return** $\boldsymbol{z}_{1:K}, \boldsymbol{\rho}_{1:C,1:K}$

#### Supplementary Methods

##### Benchmarking by using ROC curve

For each dataset with ground truth,

- Step1: Extract modified expression or posterior probability in cell-by-gene matrix for each method, each CNA state;
- Step2: Prepare ground truth matrices of binary values (1 for CNV, 0 for Neutral) for each CNA state;
- Step3: Perform ROC and AUC analysis.

To perform the benchmarking, we first converted the output of each tool to a cell-by-gene signal matrix, whose values are either modified expression (normalized/smoothed) or CNA event posterior probability indicating the presence of CNAs. Specifically, the gene scale output was directly used (XClone, inferCNV and CopyKAT) and the large genomic region scale output was mapped to the gene scale for comparison (CaSpER and Numbat). Genes overlapping with more than one genomic region would be discarded for further benchmarking analysis. Only consensus cells and genes of the 5 tools were used for each tool to extract the signal matrix for a fair evaluation.

Specifically, consensus cells were obtained by intersecting the cells of the five tools' output. Consensus genes were obtained by the same intersection way of four tools without Numbat, because Numbat only reports the posterior probabilities in the CNA regions it detects but no other regions. In this way, the consensus cells-genes matrix of Numbat is constructed by using the probabilities for reported CNA regions overlapped with the consensus gene set while setting the probabilities for other genes to be zero.

Next, the same dimensional binary ground truth matrix was built, which indicates the existence of CNA in the consensus cells and genes obtained above. The ground truth matrices are prepared for each state (copy gain, copy loss, and copy neutral LOH) independently. The number of consensus cells and genes used in BCH869 benchmarking are listed in Supp. Table S3.

Finally, ROC analysis (with True positive rate (TPR) versus False positive rate (FPR) by varying the ranking scores) was conducted to quantify the accuracy of the identification at each CNA state for each tool. Here, the signal matrix (i.e., the modified expression, for InferCNV, CopyKAT and CaSpER) and the posterior probability matrix (XClone and Numbat) are used as scores and the binary matrix from the ground truth is used as labels for ROC curve plotting. Then the area under the ROC curve (AUC) is further calculated for as a quantitative score for each tool.

See https://github.com/Rongtingting/CNV_calling_Benchmark for complete scripts for all four methods benchmarking on BCH869 dataset.

##### Details of Benchmarking on BCH869 with ground truth

BCH869 dataset is a public dataset[1], and the raw FASTQ files were generated on the SMART-seq2 platform. The FASTQ files of 492 single cells were aligned to GRCh37 (hg19) by STAR v2.7.7a.

- CNA ground truth
  Filbin et al detected 4 CNA clones that contain 489 cells. The remaining three cells are normal cells. The cell IDs of each CNA clone are obtained from the authors. The CNA ground truth for copy gain, copy loss, and LOH is publicly available in Table S7 of the original paper[1] and also presented in Fig. 1b, which were inferred from SMART-Seq2 scRNA-seq data. The complete CNA profile as ground truth is in Supp. Table S2 and also in https://github.com/Rongtingting/CNV_calling_Benchmark/blob/main/scripts/BCH869/scRNA_evaluate/data/BCH869.cnv.ground.truth.clean.0316.tsv

- Run each tool on BCH869 scRNA-seq dataset

For all five tools, 3 normal cells were used as reference cells.

For XClone (v.0.3.4), the BAF and RDR signals were generated by xcltk (v.0.1.15) preprocessing pipeline. For CNA detection, the default parameters for Smart-seq were used.

For InferCNV (v.1.8.0), the following parameters were used to detect CNV:

```
cutoff=1, cluster_by_groups=TRUE, denoise=TRUE, HMM=TRUE.
```

For CopyKAT (v.1.0.4), the following parameters were used to detect CNV:

```
id.type="S", ngene.chr=5, win.size=25, KS.cut=0.15,
distance="euclidean", n.cores=20.
```

For CaSpER (v.0.2.0), the BAF signals were generated following its manual. For CNA detection, the following parameters were used:

```
sequencing.type="single-cell", cnv.scale=3, loh.scale=3,
expr.cutoff=1, matrix.type="normalized", method="iterative".
```

For Numbat (v.1.2.1), the BAF signals were generated following its manual. The cellranger count matrix was used directly as RDR signals. For CNA detection, the following parameters were used:

```
genome="hg19", t=1e-5, gamma=5, ncores=10, plot=TRUE.
```

##### Details of Benchmarking on ATC sample

- Run each tool on ATC2 scRNA-seq dataset

For XClone (v.0.3.4), the BAF and RDR signals were generated by xcltk (v.0.1.15) preprocessing pipeline. For CNA detection, the default parameters for 10x scRNA-seq were used.

For InferCNV (v.1.8.0), the following parameters were used to detect CNV:

```
cutoff=0.1, cluster_by_groups=TRUE, denoise=TRUE, HMM=TRUE.
```

For CopyKAT (v.1.0.4), the following parameters were used to detect CNV:

```
id.type="S", ngene.chr=5, win.size=25, KS.cut=0.1,
distance="euclidean", n.cores=20.
```

For CaSpER (v.0.2.0), the BAF signals were generated following its manual. For CNA detection, the following parameters were used:

```
sequencing.type="single-cell", cnv.scale=3, loh.scale=3,
expr.cutoff=0.1, filter="median", matrix.type="normalized",
method="iterative".
```

For Numbat (v.1.2.1), the BAF signals were generated following its manual. The cellranger count matrix was used directly as RDR signals. For CNA detection, the following parameters were used:

```
genome="hg38", t=1e-5, ncores=4, plot=TRUE.
```

##### Details of Benchmarking on TNBC sample

- Run each tool on TNBC1 scRNA-seq dataset

  For XClone (v.0.3.4), the BAF and RDR signals were generated by xcltk (v.0.1.15) preprocessing pipeline. For CNA detection, the default parameters for 10x scRNA-seq were used.

  For InferCNV (v.1.8.0), the following parameters were used to detect CNV:

```
cutoff=0.1, cluster_by_groups=TRUE, denoise=TRUE, HMM=TRUE.
```

  For CopyKAT (v.1.0.4), the following parameters were used to detect CNV:

```
id.type="S", ngene.chr=5, win.size=25, KS.cut=0.15,
distance="euclidean", n.cores=20.
```

  For CaSpER (v.0.2.0), the BAF signals were generated following its manual. For CNA detection, the following parameters were used:

```
sequencing.type="single-cell", cnv.scale=3, loh.scale=3,
expr.cutoff=0.1, filter="median", matrix.type="normalized",
method="iterative".
```

  For Numbat (v.1.2.1), the BAF signals were generated following its manual. The cellranger count matrix was used directly as RDR signals. For CNA detection, the following parameters were used:

```
genome="hg38", t=1e-5, gamma=20, ncores=10, plot=TRUE.
```

##### Details of Benchmarking on GBM sample with ground truth

GBM-10x dataset is a public dataset[5], and the raw FASTQ files were generated on the 10x Genomics platform. The FASTQ files of 4416 single cells were aligned to GRCh38 (hg38) by CellRanger v.7.1.0.

- CNA ground truth

  The cell IDs of each CNA clone are obtained from the authors (personal communication). The complete CNA profile is available at https://github.com/Rongtingting/CNV_calling_Benchmark/blob/main/scripts/GBM_10x/scRNA_evaluate/data/GBM_10xscrna.celltype.cnv.agg_cnvtype.sort.tsv

- Run each tool on GBM scRNA-seq dataset

  For XClone (v.0.3.4), the BAF and RDR signals were generated by xcltk (v.0.1.15) preprocessing pipeline. For CNA detection, the default parameters for 10x scRNA-seq were used.

  For InferCNV (v.1.8.0), the following parameters were used to detect CNV:

```
cutoff=0.1, cluster_by_groups=TRUE, denoise=TRUE, HMM=TRUE.
```

  For CopyKAT (v.1.0.4), the following parameters were used to detect CNV:

```
id.type="S", ngene.chr=5, win.size=25, KS.cut=0.1,
distance="euclidean", n.cores=20.
```

  For CaSpER (v.0.2.0), the BAF signals were generated following its manual. For CNA detection, the following parameters were used:

```
sequencing.type="single-cell", cnv.scale=3, loh.scale=3,
expr.cutoff=0.1, filter="median", matrix.type="normalized",
method="iterative".
```

For Numbat (v.1.2.1), the BAF signals were generated following its manual. The cellranger count matrix was used directly as RDR signals. For CNA detection, the following parameters were used:

```
genome="hg38", t=1e-5, gamma=20, ncores=10, plot=TRUE, multi_allelic=FALSE.
```

### Supplementary Technical Notes: scCNAsimulator

#### 1 Implementation

To perform the allele-specific CNA simulation in single cells, it mainly takes an indexed BAM file and clonal CNA profile as input, and outputs a new indexed BAM file containing the desired CNA alignments. The simulator does not produce randomly generated "brand new" reads or UMIs, like scReadSim [4] did. Instead, it samples from the existing reads in the input BAM file by iterating each alignment and matching it with the clonal CNA profile.

To perform the simulations of clonal allele-specific CNAs with the input BAM file, three modules (sub-commands) are implemented, including *pileup*, *simu*, and *pipeline*. Briefly, the *pileup* module pileups allele-specific (i.e., haplotype-specific) unique molecular identifiers (UMIs) in each single cell. The *simu* module simulates CNAs based on the given clonal CNA profile and a list of haplotype-aware UMIs produced by the *pileup* module. The *pipeline* module is a wrapper that sequentially runs the *pileup* and *simu* modules. In the following section 1.1 and 1.2, we will describe the details of the *pileup* and *simu* modules.

##### 1.1 The *pileup* module pileups allele-specific UMIs

The *pileup* module is aimed to extract the haplotype-specific UMIs in single cells for each input CNA region. Currently, it relies on a list of phased bi-allelic heterozygous single nucleotide polymorphisms (SNPs) as the input source of haplotype information. These phased SNPs are typically from reference phasing, e.g., with Eagle2 [3], or generated by more sophisticated phasing methods, such as CHISEL [6] (for scDNA-seq data) and XClone [2] (for scRNA-seq data).

The *pileup* module uses multi-threading to process the input regions in parallel. In each region, the phased SNPs covered by it will be used for pileup reads from the input BAM file. For each of the covered SNP, all pileup-ed reads will be iterated and processed sequentially in the order of their start genomic positions. Processing starts with a step of quality control (QC). By default, the reads that match any of the following conditions will be filtered: (1) FLAG includes any of UNMAP, SECONDARY, QCFAIL bit, (2) aligned length<30nt, (3) mapping quality<20, (4) or being singletons. After QC, the haplotype information (i.e., the `HAP`) of the iterated read and also its source UMI can be obtained if the read contains any of the two alleles (i.e., the `REF` or `ALT`) of the phased SNP. Then, the haplotype-aware UMIs will be outputted and used by the downstream *simu* module.

Notably, in the module, (1) the phased SNPs will be discarded if their aggregated UMI counts pileup from all cells is smaller than `minCOUNT` (default is 1) or the frequency of the minor allele is smaller than `minMAF` (default is 0). (2) UMIs with conflicting haplotype information will be discarded, e.g., when its reads cover both alleles of one (phased) SNP.

##### 1.2 The *simu* module simulates clonal CNAs

The *simu* module simulates CNAs by forking or discarding UMIs based on the given clonal CNA profile, and output a new indexed BAM file. It starts by iterating each read in the order of their start genomic positions. The reads whose source cells `CELL` are not in input CNA clones `CLONE_ID` will be outputted without further processing. The key step to process the remaining reads is to obtain the allele-specific copy number `CN` of their belonging UMIs, while the procedures are different for haplotype-aware UMIs and ambiguous ones.

Specifically, we first try querying the allele-specific UMI list produced by the *pileup* module, with the cell barcode `CELL` and UMI barcode `UMI`, to obtain the allele/haplotype information `HAP` of the read and also the overlapping CNA region `REG_ID`. If the UMI is haplotype-aware (i.e., UMI barcode is in the allele-specific UMI list), then we further query the input clonal CNA profile with the `REG_ID` and `CLONE_ID` to obtain its allele-specific copy number (`CN0` or `CN1`).

Otherwise, if the source UMI of the read is ambiguous (i.e., not haplotype-aware), then we first check whether the `CN` of the UMI is available (i.e., any of its reads has been processed and `CN` recorded). If yes, then we use the recorded value directly. Otherwise, we query the CNA profile by matching the genomic range of the read with the CNA regions, to obtain the corresponding allele-specific copy numbers `CN0` and `CN1`. As the UMI is ambiguous, we randomly select from `CN0` and `CN1` with equal probabilities (i.e., 0.5 vs. 0.5), as the allele-specific copy number of the UMI and record it for later use.

Based on the allele-specific copy number `CN`, we fork or discard the reads for both haplotype-aware and ambiguous UMIs. Specifically, if `CN` is 0, we discard the read/UMI; otherwise, we fork the read/UMI `CN` times. Note that when forking UMIs, the QNAME and UMI barcode (`UB` tag) of the forked reads are modified (currently adding a suffix) to make them distinct from the original ones, hence are also unique (`CELL+UMI`) in the whole output BAM file. The QNAME and UMI barcode of other reads are also modified accordingly to make all output reads have the same format.
